# Microbiome contribution to *Indy* longevity in *Drosophila*

**DOI:** 10.64898/2026.03.25.714291

**Authors:** Danielle N. A. Lesperance, Shivani Padhi, Jacob Macro, Sara Olson, Ella Stanwood, Kavitha Kannan, Brenton Graveley, Blanka Rogina, Nichole A. Broderick

**Affiliations:** Department of Molecular and Cell Biology, University of Connecticut, Storrs, CT 06269, USA; Department of Genetics & Genome Sciences, School of Medicine, University of Connecticut Health, Farmington, CT, 06030, USA; Institute for Systems Genomics, School of Medicine, University of Connecticut Health, Farmington, CT, 06030, USA; Department of Biology, Johns Hopkins University, Baltimore, MD 21218, USA

**Keywords:** *Indy*, Microbiome, Lifespan, *Drosophila melanogaster*, Aging

## Abstract

Reduction in the *Indy (I’m not dead yet)* gene, a plasma membrane citrate transporter, in *Drosophila* and its homolog in worms extends lifespan by promoting metabolic homeostasis. *Indy* reduction delays the onset of aging-associated pathology in the fly midgut, including preservation of intestinal barrier integrity and intestinal stem cell homeostasis. Gut microbiota has broad impacts on host metabolism, health, and aging. Age-related dysbiosis impairs intestinal barrier function and drives mortality. However, the underlying mechanisms that link increased microbial load to frailty and negative effects on health remain mostly unclear. Here we show that *Indy* heterozygote flies have significantly lower bacterial load and increased diversity during aging compared to controls. However, the presence of the microbiome was not required for *Indy* lifespan extension, though removal of microbes did enhance the effects of *Indy* reduction on longevity, suggesting potential interactions between the microbiome and *Indy*. *Indy* down-regulation was linked to reduced expression of the JAK/STAT signaling ligands Upd3 and Upd2 in the midgut of young flies, which likely contributes to preserved intestinal stem cell homeostasis. Altogether, our results suggest that *Indy* reduction impacts microbiome load and composition, which preserves gut homeostasis and extends lifespan through impacts on JAK/STAT signaling pathway.

**Significance Statement:** *Indy* is a fly homologue of mammalian SLC13A5 (mSLC13A5) plasma membrane citrate transporter, a central metabolic regulator involved in health, longevity, and disease. Reduction of fly *Indy* gene activity preserves metabolic and intestinal stem cell homeostasis and extends longevity. Gut microbiota impacts host metabolism, health, and aging. Here we show that *Indy* reduction prevents age-associated increases in bacterial load and expression of the JAK/STAT signaling ligands Upd3, and Upd2, while maintaining microbiome diversity. These changes likely slow activation of epithelial cell turnover in the gut and contribute to downstream lifespan effects. As the role of INDY and microbiome are conserved across organisms, our study provides a framework to study underlying mechanisms of the effects of reduced *Indy* and the microbiome on health and longevity.

## Introduction

### *Indy* and microbiome have vital roles in health and longevity

The aging process in organisms is characterized by loss of homeostasis, accumulation of cell damage, and decline in physiologic function (1–3). Key factors that contribute to aging and its related pathologies have been identified using model organisms including mice, worms, and *Drosophila melanogaster* (3–5). Microbiota imbalance has been linked to a growing number of human disorders including obesity, cardiovascular disease, type-2-diabetes and inflammatory bowel disease. One of the major risk factors for dysbiosis is aging. Age-associated dysbiosis of the intestinal microbiota has been implicated in intestinal barrier impairment, activation of intestinal immune response and increased frailty. Increased microbial load in the fly midgut has been associated with age-related intestinal dysfunction and increased mortality (6–10).

The *Indy (I’m not dead yet)* gene encodes the fly homolog of a mammalian SLC13A5 plasma membrane citrate transporter (11–14). Reduction of *Indy* gene activity in flies or its homologs in worms, rats, and mice has beneficial effects on energy balance by affecting the levels of cytoplasmic citrate, which plays a central role in metabolism (15–21). Citrate levels affect glucose and lipid metabolism, and energy production by mitochondria. In flies, INDY protein is highly expressed in the midgut, fat body, and oenocytes, all of which are important sites of fly metabolic regulation (11–13, 22–24). *Indy* reduction alters the metabolism of flies, worms, mice and rats, in a manner akin to changes associated with caloric restriction (CR) (11, 15–17, 19, 25–28). Reduction in *Indy* gene expression extends longevity of *Indy^206^/+* heterozygous flies and in other *Ind*y heterozygous alleles, compared to control flies, delays the onset of intestinal barrier disfunction, and preserves intestinal stem cell homeostasis, and midgut integrity and function (11, 22, 23, 25, 26, 29). Maximal life span extension was observed in *Indy* heterozygous flies, and their lifespan is not further extended by CR, providing further evidence that *Indy* and longevity pathways overlap. While *Indy* homozygous flies live longer than control flies when aged on a conventional laboratory diet or a high caloric diet, their lifespan is shorter than *Indy* heterozygous flies most likely because they are malnourished (20). CR without malnutrition delayed the onset of aging associated pathologies, and extended lifespan in a variety of species (30–33). Beneficial effects of CR on the microbiome and global host metabolism have been reported (34, 35). Transplantation of the microbiome from female mice aged for 8 weeks on a very-low calorie diet to obese female mice increased recipient microbiome alpha diversity, decreased their body fat accumulation, and improved glucose tolerance compared to mice that received microbiome from ad libitum fed mice (34). In addition, antitumor effects of CR have been linked to changes in the gut microbiome (36). However, the level of CR is important for the effects on microbiome, as severe CR improved metabolic health in a randomized human interventional study, but led to a decrease in bacterial abundance and altered microbiome composition (37). Transplantation of the post-CR diet microbiome decreased adiposity and body weight of recipient mice by impairing nutrient absorption and enriching for pathobionts including, *Clostridioides difficile*. In another randomized controlled study, CR was shown to affect microbiome composition and shift plasma metabolites to a longevity-related metabolic pathway signature (38). These studies highlight the importance of interactions between nutrients and microbiota and their effects on metabolism.

The JAK/STAT signaling pathway has a key role in epithelium renewal (6, 39–41). *Drosophila* cytokines Upd3, Upd2 and Upd are released from different cells type in the intestine to promote stem cell proliferation through activation of the JAK/STAT pathway. While JAK/STAT is important for initiating damage repair pathways in response to stress, sustained activation of JAK/STAT, as occurs in old age, disrupts homeostatic processes of the gut, contributing to organismal death (42). These age-related increases in the JAK/STAT signaling are attributed in large part to increased microbiome density, a commonly reported phenotype in aging *D. melanogaster* (7, 8, 10, 41, 43). Overexpression of the JAK/STAT pathway leads to increased enterocyte damage and aberrant ISC proliferation resulting in midgut hyperplasia and reduced lifespan. Age-related increases in gut damage and microbiome density are associated with higher ROS production further contributing to activation of the JAK/STAT pathway. Previously, we reported that the midgut of *Indy* flies have reduced ROS accumulation and increased expression of genes encoding ROS-detoxification enzymes, which reduces age-associated hyperproliferation of ISCs (23, 24). Given the impact of the microbiome on gut damage in aging and JAK/STAT activity, it is possible that the beneficial effects of *Indy* reduction are mediated through impacts on the fly microbiome and reduced JAK/STAT signaling pathway activation. Therefore, we hypothesized that reduction in *Indy* gene activity could also alter the gut microbiome, which in turns reduces JAK/STAT signaling. As such, reduced *Indy* gene expression would lead to lower bacterial load during aging, translating to reduced JAK/STAT signaling, which would be consistent with the previously reported preservation of intestinal homeostasis into old age, and lifespan extension in heterozygous flies (22, 23).

To address our hypothesis, we characterized the microbiomes of young and old control and *Indy^206^/+* flies and found significant decrease in bacterial load and increased bacterial diversity in aging *Indy^206^/+* flies compared to age-matched control flies. Additionally, we compared lifespan of control and *Indy^206^/*+ flies with and without a microbiome to determine if lifespan extension in *Indy* heterozygotes was dependent on the presence of the microbiome. We found that the presence of the microbiome was not required for *Indy^206^/+* lifespan extension, but removal of microbes did enhance the effects of *Indy* reduction on longevity. In addition, transcriptomic analysis on the midgut of control *yw* and *Indy* on conventional and AX conditions revealed reduction in expression of the JAK/STAT signaling ligand Upd3, and Upd2 in young *Indy^206^/+* flies, which leads to slower activation of epithelial cell turnover in the gut and most likely contributes to preserved intestinal stem cell homeostasis of *Indy* flies. Taken together, our study demonstrated that reduced *Indy* impacts microbiome load and diversity, which likely has downstream effects on gut homeostasis and aging phenotypes.

## Results

### Presence of the microbiome augments but is not required for lifespan extension in *Indy* heterozygotes

To determine if the previously reported differences in lifespan between control and *Indy^206^/+* flies depend on the presence of the microbiome, we assessed lifespan of both conventional (containing an un-altered microbiome) and axenic (containing no microbes) flies. Flies were passaged twice per week. Median lifespan of *Indy^206^/+* exceeded that of controls by 12 days in males (21%) and 8 days in females (21%) in conventional conditions (Fig. 1A,B, Table S1). In axenic conditions, median lifespan of *Indy^206^/+* flies was 7 days (10%) longer than controls in males and 16 days (50%) longer in females (Fig. 1C,D, Table S1, Fig. S1A-D), suggesting that removal of bacteria impacted control and *Indy/+* flies to different extents. Thus, while the microbiome is not necessary for *Indy^206^/+* lifespan extension, microbes may impact control and *Indy* flies in different ways, potentially indicating direct interactions between the microbiome and *Indy* and/or citrate metabolism. While both male and female *Indy^206^/*+ and male *yw* live longer in axenic condition compared to conventional, control female lifespan was not significantly different in axenic conditions (Fig. S1C, Table S3). Lifespan studies were repeated by passing flies daily. Median lifespans of *Indy^206^/+* male flies were significantly increased compared to *yw* control male flies in conventional and in AX condition, while female *Indy^206^/+* flies showed small changes in median lifespan in conventional condition, and a significant increase in median lifespan in AX conditions compared to *yw* control females (Fig. 1E,F, Fig. S1E-H, Table S3). Consistent with lifespans of flies that were passaged twice per week, males and female *Indy^206^/+,* and *yw* male lived longer in AX conditions, while *yw* female lived shorter in AX condition compared to aging on a conventional diet (Fig. 1 E,F, Table S4).

**Figure 1.**
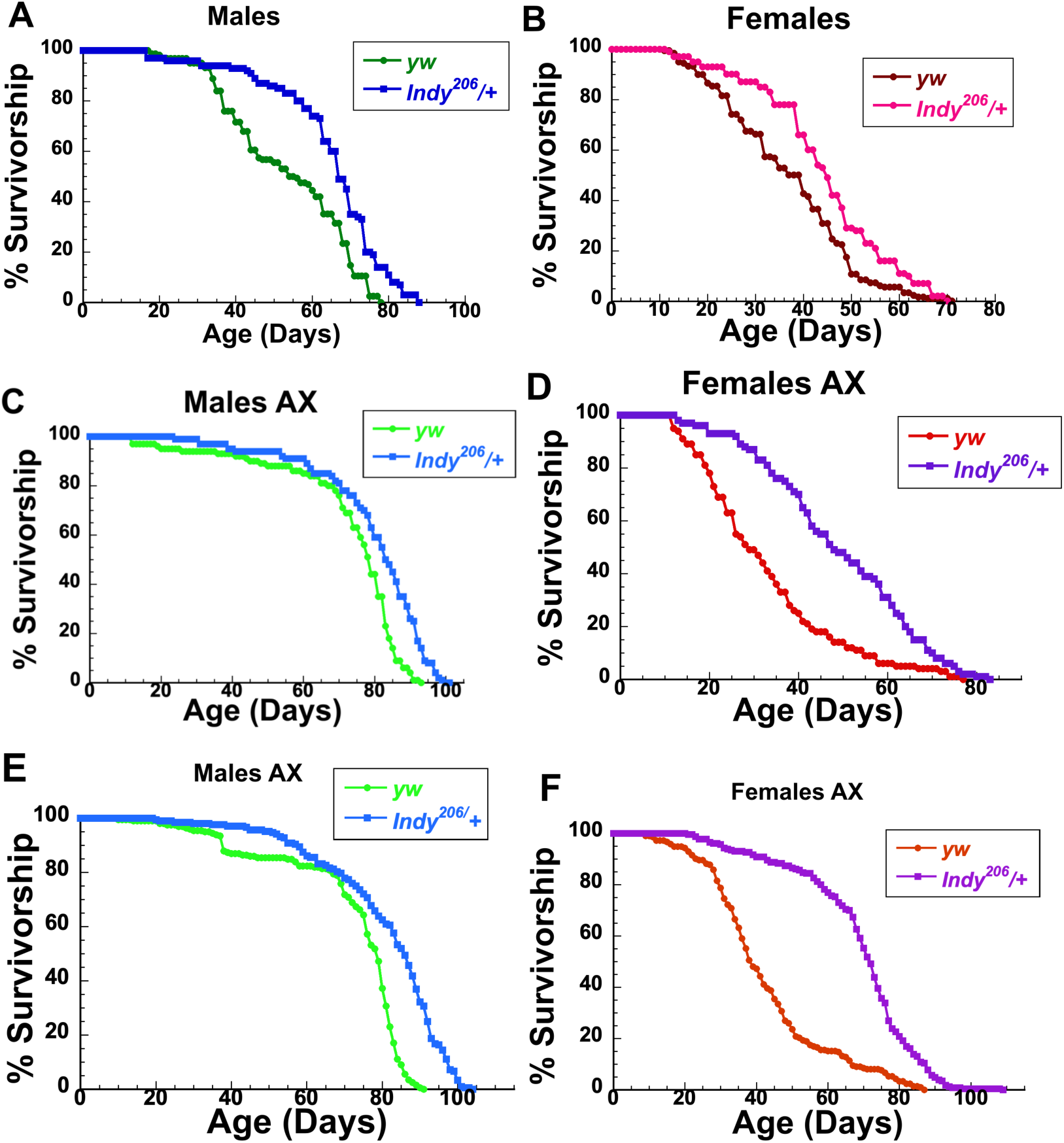
A-D) Lifespan of conventional and axenic *yw* control and *Indy^206^/+* heterozygous flies passed twice per week. Longevity of conventional (A-B) and axenic (C-D) *yw* control (green (A), brown (B) light green (C), orange (D)) and *Indy^206^/+* (blue (A,C), magenta (B), purple (D)) male (A,C) and female (B,D) flies passed twice per week expressed as Kaplan-Meier survival curves. N= 131-187 individual flies per condition spread across 10 vials that were passed twice weekly. **E-F) Effects of axenic culture on lifespan of *yw* control and *Indy^206^/+* flies passed daily.** Longevity of axenic *yw* control ((light green (E), orange (F)) and *Indy^206^/+* ((blue (E), purple (F)) male (E) and female (F) flies passaged daily expressed as Kaplan-Meier survival curves. AX: axenic.

### Bacterial load is lower in aging *Indy^206^/+* compared to control flies

It has been reported that aging-associated gut pathology is exacerbated by increased bacterial load in aging flies (7, 40, 43). To determine the extent to which delayed aging phenotypes observed in *Indy^206^/+* flies may be attributed to microbiome load, we performed culture-dependent microbiome analysis of young and old *yw* controls and *Indy^206^/+* flies via plating of whole fly homogenates on MRS agar (Fig. 2A). We found that while bacterial load was similar in the two genotypes at 7 days old, 40-day old *Indy^206^/+* flies had a roughly 10-fold lower microbiome load compared to their *yw* counterparts (Fig. 2C). Repeated analysis showed that the total bacterial load was lower in *Indy^206^/+* female flies at both 7 and 40 days (Fig. S2A). In addition, we observed some differences in microbiome composition between the two genotypes, especially at 40 days old, as *Indy^206^/+* flies had a higher abundance of a specific isolate (isolate 2) that we determined by 16S rRNA sequencing (V3/V4 region) was most closely related to *Acetobacter pomorum* or *pasteurianus* (Fig. 2D). Both genotypes contained similar levels of another isolate, closely related to either *A. malorum* or *A. orleanensis* (Fig. 2D).

**Figure 2.**
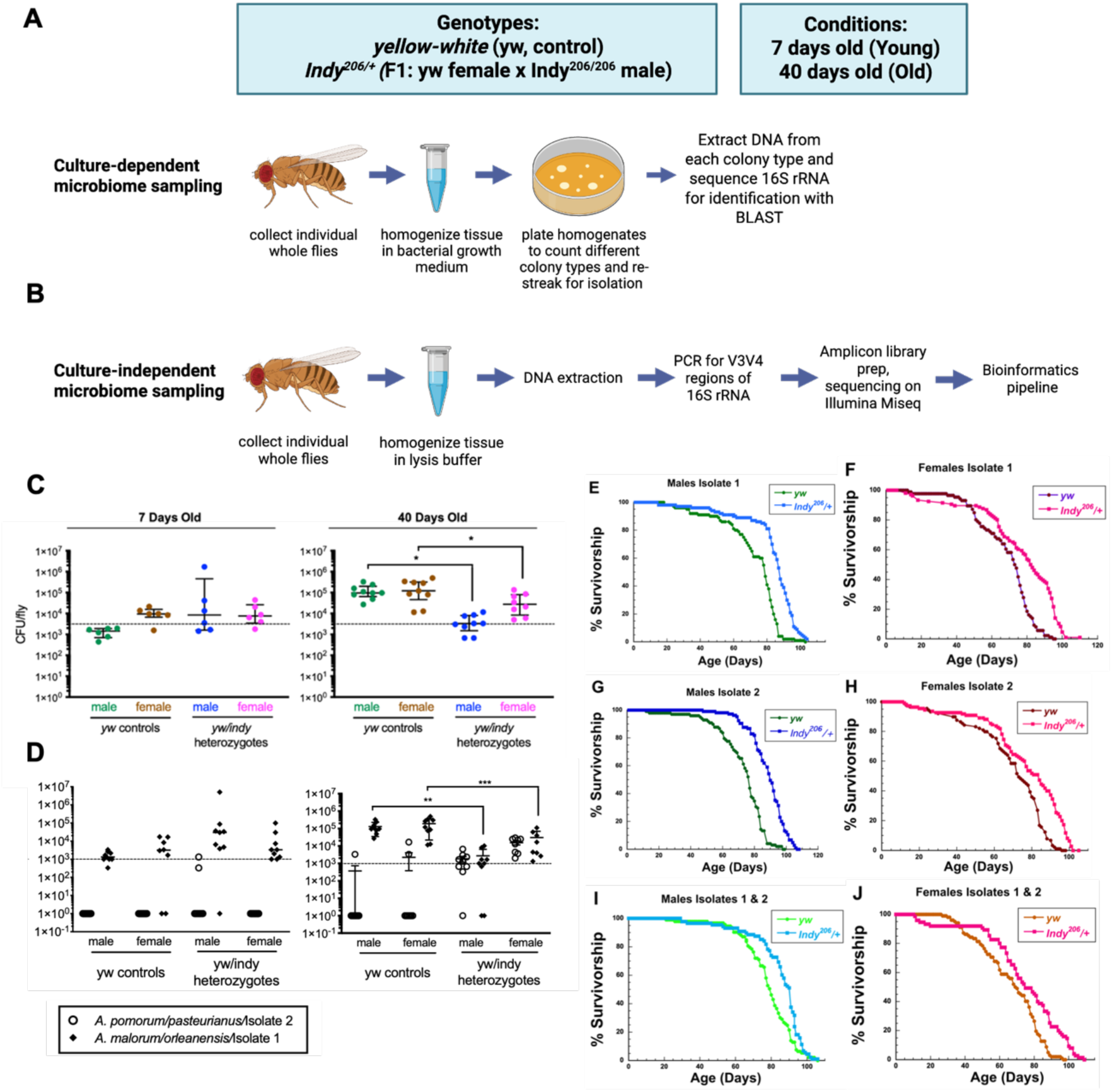
Experimental diagram: Workflow of culture-dependent (A) and culture-independent (B) microbial sequencing from individual whole flies, which were homogenized in bacterial growth media, plated and extracted DNA was sequenced (A), or DNA was extracted from homogenate, PCR for V4 region of 16S rRNA and sequenced (B). **C) Total bacterial load in 7 day and 40-day old *yw* controls and *Indy^206^/+* flies.** Bacterial counts are expressed as colony forming units (CFU) per fly, as determined by growth of colonies on MRS agar. Points represent individual flies; lines and error bars represent median and interquartile range of data points. **D) Load of bacterial isolates 1 and 2 detected in 7 day and 40 day old *yw* controls and *Indy^206^/+* flies.** Bacterial counts are expressed as colony forming units (CFU) per fly, as determined by growth of colonies on MRS agar. **C, D** Points represent individual flies; lines and error bars represent median and interquartile range of data points. Statistical differences between *yw* controls and *Indy^206^/+* CFU counts were determined via two-way ANOVA with Bonferroni multiple comparisons tests. N= 3 replicate experiments collecting 3 individual flies per treatment group (n=9 flies per genotype/age). Statistical results not shown were non-significant. Significance is expressed as follows: ns, not significant (*P* > 0.05); *, *P* ≤ 0.05; **, *P* ≤ 0.01; ***, *P* ≤ 0.001; ****, *P* ≤ 0.0001. **E-J) Lifespan of gnotobiotic *yw* control and *Indy^206^/+* heterozygous flies.** Longevity of gnotobiotic control *yw* and *Indy^206^/+* male (E,G,I) and female (F,H,J) flies containing Isolate 1 (E,F), Isolate 2 (G,H), or a mix of both isolates (I,J) expressed as Kaplan-Meier survival curves. Statistical differences between control and heterozygote lifespan were determined via Log-rank analysis. N= 84-125 individual flies per condition spread across 7-13 vials passed twice weekly. All survivorships of *Indy^206^/+* were significantly longer compared to *yw* control flies P::0.0001=****. het: heterozygote *Indy^206^/+*

### Lifespan extension of *Indy^206^/+* flies is not dependent on presence of *Acetobacter spp*

To determine if the presence of either cultured *Acetobacter* isolate was responsible for the lifespan differences between conventional control and *Indy^206^/+* flies, we generated gnotobiotic flies by inoculating axenic embryos with isolate 1 (*A. pomorum/pasteurianus*), isolate 2 (*A. malorum/orleanensis*), or a mix of both. We recorded lifespan of each gnotobiotic condition and found that, regardless of bacterial treatment, *Indy^206^/+* lifespan exceeded that of controls as seen in both conventional and axenic conditions (Fig. 2E-J). These results suggest either that: 1) neither *Acetobacter* isolate impacts lifespan extension of *Indy^206^/+ flies*, or 2) that both isolates impact longevity in a similar fashion.

### Microbiome composition differs between control and *Indy* flies

To further investigate microbial community composition in each genotype, we performed culture-independent 16S rRNA microbiome sequencing on *yw* controls, *Indy^206^/Indy^206^* homozygous, and *Indy^206^/+* heterozygous flies (Fig. 2B). While this analysis could not distinguish between species of *Acetobacter* as in Fig. 2, it did reveal the presence of a member or members of the *Lactobacillus* genus, which are common members of the *D. melanogaster* microbiome. In both male and female flies, *Lactobacillus* was most prevalent in *Indy^206^/Indy^206^*, and *Indy^206^/+* flies, and was only present in a few control flies, particularly at 7 days old (Fig. 3A). By 40 days, most *Indy^206^/Indy^206^*, and *Indy^206^/+* flies were still associated with *Lactobacillus spp*., whereas it was mostly absent from *yw* aged-matched controls (Fig. 3A). This difference in *Lactobacillus* presence between genotypes was reflected in the beta and alpha diversity metrics, which show greater differences between *Indy^206^/+* and *yw* controls than between *Indy* heterozygous and homozygous flies (Fig. 3B-E). Furthermore, *Indy^206^/Indy^206^*and *Indy^206^/+* male and female flies have increased Shannon diversity compared to control yw flies (Fig. 3 F,G). A similar increase in Shannon diversity was confirmed in young and old *Indy^206^/+* female flies compared to controls in a second set of experiments (Fig. S2B). Along with our culture-dependent analysis, these differences in microbial diversity between aging *yw* controls and *Indy^206^/+* suggest that microbial diversity, in addition to total microbial load, may be impacting aging.

**Figure 3.**
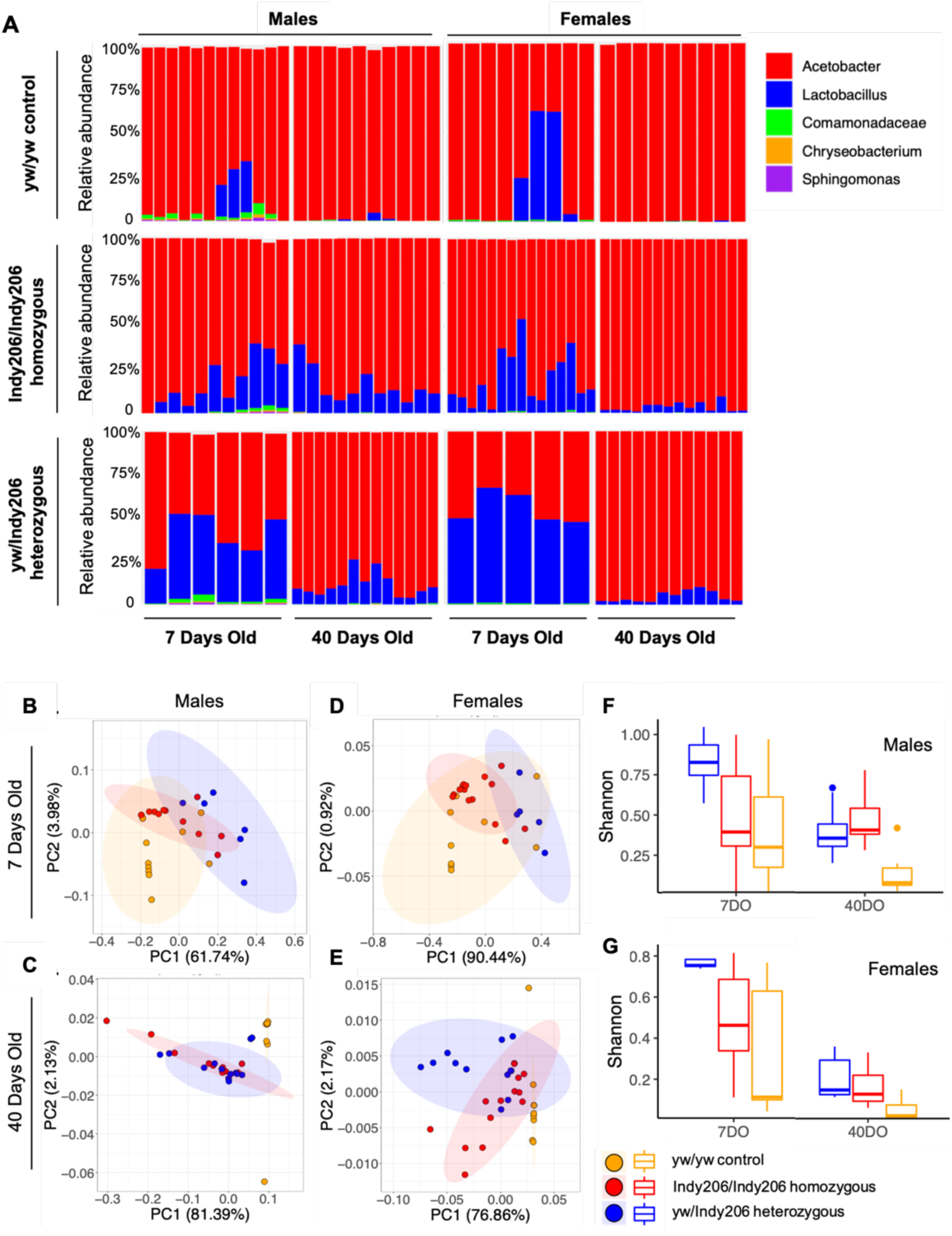
A) Microbial community composition of young and old *yw* control, *Indy^206^/Indy^206^*, and *Indy^206^/+* flies. Plots represent relative abundance of top five genera of bacteria present in individual whole flies. B-G) Beta and alpha diversity of young and old *yw* control, *Indy^206^/Indy^206^*, and *Indy^206^/+* flies. (A-D) Principal component analysis (PCoA) of Bray-Curtis distances between microbiomes of individual fly samples. Genotypes are indicated by different colors. Shaded ellipses represent 95% confidence intervals for clusters of each group. **(F,G) Shannon diversity** of each condition separated by males (F) and females (G). Boxes range from the 25^th^ to 75^th^ percentile values for individual samples within each group; middle line shows the median; whiskers represent minimum and maximum values; individual points outside of minimum and maximum show outliers.

### *Indy* reduction has a significant impact on host transcriptome

To determine molecular mechanisms and pathways underlying longevity extension and the effects of the reduced microbiome load in *Indy^206^/+* flies, we performed RNA-Seq analysis. We determined RNA-Seq profiles using three biological replicates at each time point and of each genotype. Bulk RNA was extracted from dissected midguts of female *Indy^206^/+* and *yw* control flies at 7 and 40 days of age, reared under conventional or axenic conditions. A sample-to-sample distance heatmap shows genotype and age to be the driving force for distance, indicating distinct transcriptional profiles across age, genotypes and rearing condition (Fig. S3A). This is corroborated by transcriptome-wide principal component (PC) analysis, which indicates that biological samples from the same age and genotype cluster together and are clearly distinct (Fig. 4A). PC analyses of samples from the AX and conventional reared flies are more similar at age 7 and partially overlap, while samples of AX and conventional reared flies at 40 days are more distant (Fig. 4A).

**Figure 4:**
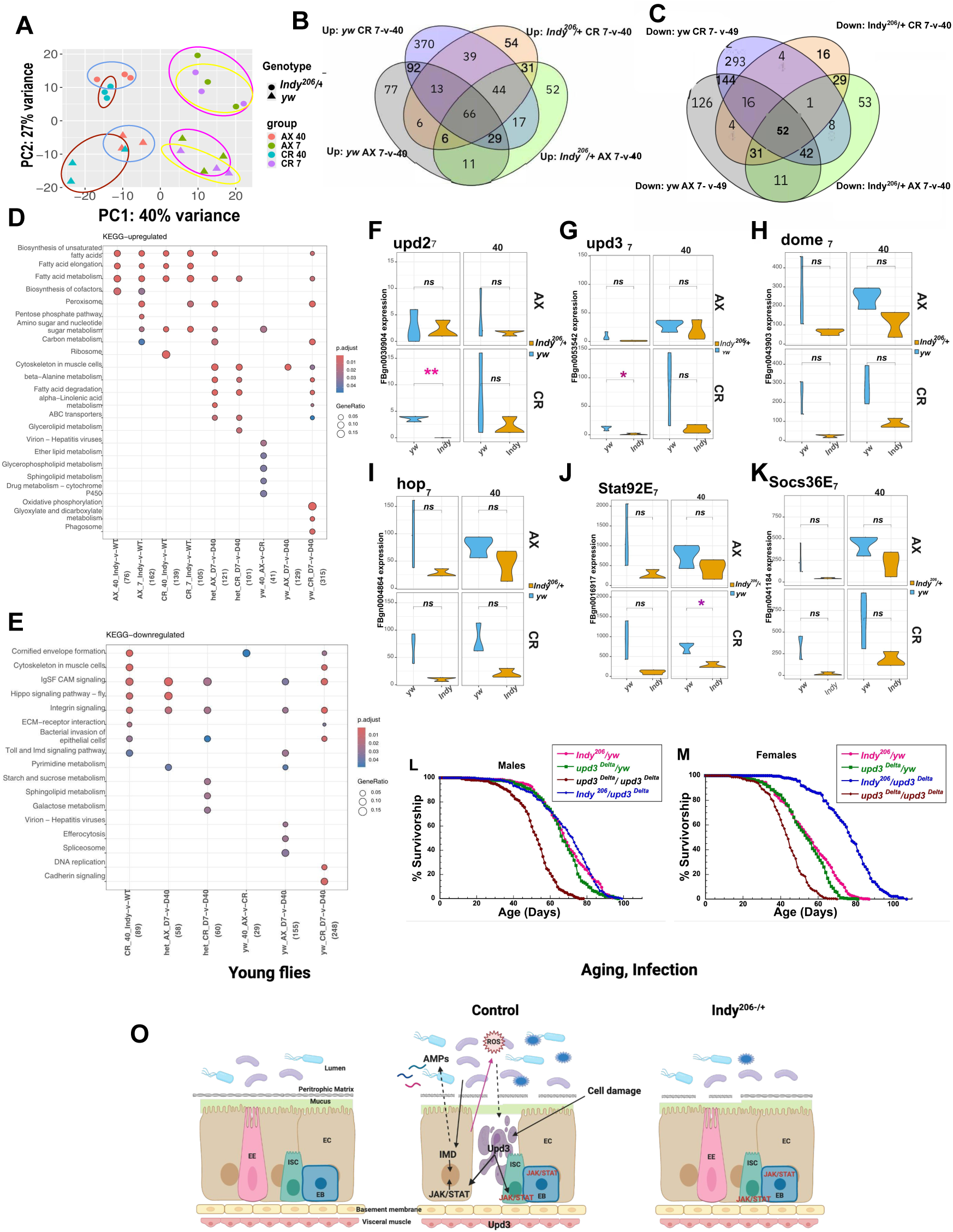
Effects of *Indy* reduction, age and AX condition on midgut transcriptome. **A)** PCA plots of differentially expressed genes in midgut samples from *Indy^206^/+* and *yw* control flies in conventional (CR) and axenic (AX) condition at 7 and 40 days of age. B,C) Venn diagrams show common or unique upregulated (Up) (B) or downegulated (Down) (C) DE genes between *yw* control and *Indy^206^/+* in flies reared in AX or conventional conditions: Up: yw CR 7-v-40, Up: yw AX 7-v-40, UP: *Indy^206^/+* CR 7-v-D-40 and Up: *Indy^206^/+* AX 7-v-40. C) The same comparison shown for downregulated DE genes. FDR threshold: 0.02, logFC threshold: 0 D,E) Functional enrichment analysis shows KEGG pathways of upregulated (D) or downregulated (E) DE genes associated with *Indy* reduction, aging, conventional vs AX conditions compared to *yw* controls. logFC threshold: 0.2. F-K) ***Indy* reduction affects JAK/STAT signaling pathway**. *Indy^206^/+* and *yw* (control) differentially expressed gene analysis of count data of Upd2 (F), Upd3 (G), dome (H), hop (I), Stat92E (J) and Socs36E (K) at 7 and 40 days of age in conventional (CR) and AX conditions. L, M) The epistatic relationship between the *Indy* and JAK/STAT longevity pathways. Survivorships of male (L) and female (M) *upd3^Delta^/upd3^Delta^*(brown), *upd3^Delta^*/+ (green), *Indy^206^/+* (magenta)*, and upd3^Delta^*/+;*Indy^206^/+* (black) flies expressed as Kaplan-Meier survival curves. Survivorship were determined by log-rank analysis using JMP16 program. N= 253-293 individual flies per genotypes across 12 vials that were passed daily. N) Proposed model of beneficial effects of *Indy* reduction on intestinal stem cell homeostasis. Intestinal integrity is preserved in young and aging *Indy* flies, compared to aging control flies. *Indy* reduction decreases bacterial load, increases microbial diversity and decreases JAK/STAT signaling pathway including Upd3 levels. *Indy* flies have also decreased ROS production in the midgut that all lead to longer lifespan.

### Difference in age-associated DE between of control *yw* and *Indy* on conventional and AX conditions

The generation of axenic fly lines eliminates both gut and substrate specific microbes, as well as their compounds, therefore, our study can distinguish between effects of microbiome and *Indy* reduction on transcriptome of the midgut. To search for common upregulated and downregulated age-associated DE genes, we compared *yw* control and *Indy^206^/+* age-specific DE genes in flies reared in AX and conventional conditions (CR): Up:*yw* CR 7-v-40, Up:*yw* AX 7-v-40, Up: *Indy^206^/+* CR 7-v-40 and Up: *Indy^206^/+* AX 7-v-40 (Fig.4B). Similar comparison was performed for downregulated common and unique DE (Fig. 4C). A Venn diagram shows only 66 commonly upregulated and 52 commonly downregulated DE genes across control and *Indy^206^/+* flies in AX and conventional conditions (FDR 0.02 (Fig. 4B,C). Common upregulated genes belong to four KEGG pathways including metabolic pathways, carbon metabolism, cytoskeleton and motor proteins. Highest number of unique upregulated and downregulated DE genes are observed in *yw* CR 7-v-40 females N=370, and N=293, respectively (Fig. 4B,C). This illustrates that *yw* flies experienced the largest age-associated changes when exposed to microbiome while reared in conventional conditions. As a consequence of increased bacterial load, there is an upregulation of the members of Cytochrome P450 family including Cytochrome P4504g1, Cyp6a22, Cyp12d1-p, Cyp12d1-d. The number of age-associated DE genes is much lower in both AX and conventional *Indy^206^/+* flies, consistent with lower bacterial load observed in aging *Indy^206^/+* flies (Fig. 4B,C). There are 31 common upregulated DE genes in *Indy^206^/+* in AX and conventional conditions, that include Glutathione S transferase E10, a member of the glutathione pathway, and the Phosphoenolpyruvate carboxykinase 2 (Pepck2), a mitochondrial enzyme that is the key regulator in gluconeogenesis. Common downregulated DE genes in *Indy^206^/+* AX and conventional aging flies includes tobi, target of brain insulin.

KEGG pathway analysis revealed upregulated fatty acid (FA) metabolism in samples from *Indy^206^/+* flies when compared with controls in AX and conventional conditions at 7 or 40 days of age (Fig. 4D). Upregulated biological processes include FA biosynthetic processes, FA elongation, peroxisome, pentose phosphate pathway and ribosome (Fig. 4D). FA metabolism is upregulated in *Indy* flies due to the shift to using FA oxidation as the preference for substrate oxidation. Similar changes in fat metabolism were reported in calorie restricted flies and rodents (30, 44). Comparison of the transcriptome of 7-day old *Indy^206^/+* flies to 40-day old *Indy^206^/+* kept in both AX and conventional conditions, revealed similar upregulation of carbon metabolic processes and FA metabolic processes as *Indy* flies age (Fig. 4E). Hippo signaling pathway is downregulated in 40 days old *Indy^206^/+* flies compared to *yw* control flies in CR conditions, which includes downregulation of cell proliferation, consistent with our previous report that *Indy* flies have reduced age-associated increase in intestinal stem cell (ISC) proliferation (23). There is also downregulation of Hippo signaling pathway in *Indy* flies when 7 and 40 days old flies aged in AX condition were compared. In addition, *Indy* flies have downregulated bacterial invasion of epithelial cells, as well as Toll and Imd signaling pathways, in agreement with reduced bacterial load observed in 40 days old *Indy* flies (Fig. 4E).

### The role of the JAK/STAT signaling pathway in *Indy* longevity

We next examined if there is a link between reduced *Indy* expression and the JAK/STAT signaling pathway. We determined the levels of expression of Upd3, and Upd2, which are released from different cells type and promote proliferation by activation of the JAK/STAT pathway and epithelial cell turnover in the gut. We found significant decrease in the expression levels of Upd2 and Upd3 in *Indy^206^/+* flies aged in a conventional condition, but not in flies kept in AX condition at 7 days of age (Fig. 4F,G). We also determined expression levels of other members of JAK/STAT signaling (Hop, Stat92E and Socs36E) in the midguts of *Indy* and control flies. Hop is *Drosophila* homologue of mammalian JAK, Stat92E is *Drosophila* homologue of STAT transcriptional factor and Socs36E is suppressor of cytokine signaling. There is a significant decrease in the Stat92E levels of expression levels at age 40 in CR condition (Fig. 4J). No differences were found in levels of Upd2, Upd3, Hop, and Socs36E at age 40 days in females that were kept in either conventional or axenic conditions (Fig. 4H-K). The levels of Upd2 and Upd3 were very low in *Indy^206^/+* flies compared to control flies at 7 days of age kept in conventional condition consistent with lower bacterial load in *Indy* flies at 7 days (bacterial load Fig. 2C). While the levels of expression of *Dome* and *hop* in *Indy* flies were lower compared to control flies, the levels were not significantly different in females reared on either conventional condition or AX conditions (Fig. 4H,I). Heatmap illustrate trend of lower expression levels of Upd2, Upd3, Hop, Dome, Socs36E and Stat92E in all *Indy^206^/+* samples compared to *yw* control with exception of *yw* samples isolated from 7 days old female in AX condition. These results are consistent with the lack of a microbiome in *yw* AX females and lower microbial count in *Indy^206^/+* flies (Fig. S3B).

We identified specific changes in the levels of immune and stress response genes in the midgut of control and *Indy* flies reared under conventional and axenic conditions. In aged flies, intestinal barrier dysfunction is associated with increased expression of immunity-related genes such as antimicrobial peptides (AMPs). To examine the role of the immune response in preserved intestinal integrity of *Indy* flies we determined expression levels of AMPs responsive to the microbiome and with activity predominately against Gram-negative bacteria (Diptericin A (Dpt A), Diptericin B (Dpt B)), as well as Drosocin (Dro), which is mostly active towards Gram-positive bacteria, and Drosomycin (Drs)), an antifungal peptide, using mRNA isolated from *yw* and *Indy^206^/+* females. There was no significant difference in levels of mRNA between *Indy* and control samples, however heatmap illustrates a trend of lower levels of AMPs genes in *Indy^206^/+* flies (Fig. S3C-G).

### The role of Upd3 in *Indy* longevity extension

To determine if *Indy* and JAK/STAT longevity pathways overlap we examined the epistatic relationship between these two pathways. Reducing *Indy* or JAK/STAT signaling extends fly lifespan, while overexpressing JAK/STAT activity reduces lifespan. If *Indy* and JAK/STAT pathways overlap, longevity of flies with simultaneous *Indy* and JAK/STAT reduction will be similar to longevity of flies with reduction in only *Indy* or Upd3. In contrast, if the two pathways are independent, longevity of flies with simultaneous reduction in both genes will be longer compared to a single mutant. To examine the role of Upd3 in longevity extension of *Indy* flies, we determined lifespan of *Indy^206^/+*, *upd3^D^*/+ heterozygous and *upd3^Delta^*/ *upd3^Delta^* homozygous flies, and flies double heterozygous for *Indy^206^/+* and *upd3^Delta^*/+. *upd3^Delta^*/+ allele has a 13,464 bl deletion that removes the first three exons of Upd3 (including the start ATG) made by imprecise excision of P{XP}upd3[d00871] and is shown to lack function (45). Both male and female *upd3^Delta^/upd3^Delta^*homozygous flies have the shortest median and maximal lifespan. *upd3^Delta^*/+ male and female flies have 26% and 23% longer median lifespan compared to homozygous flies respectively (Fig. 4L,M, Table S5). Median longevity of *Indy^206^/+* flies is similar to *upd3^Delta^*/+ flies when compared to homozygous *upd3^Delta^* flies, (Median lifespan: males live 38%, females 27% longer.) However, *upd3^Delta^*/+;*Indy^206^/+* male flies have a median lifespan 38% longer, while female flies live 77% longer compared to homozygous *upd3^Delta^* flies (Table S5). These finding suggest that mechanisms underlying longevity extension observed in *Indy^206^/+* and *upd3^Delta^*/+ partially overlap, but also have independent effects on longevity, more notably in females.

## Discussion

### INDY and microbiome have vital roles in health and longevity

Microbiome imbalance has been linked to number of human disorders including obesity, cardiovascular disease, type-2-diabetes and inflammatory bowel disease. One of the major risk factors for dysbiosis is aging. Age-associated dysbiosis of the intestinal microbiota has been implicated in intestinal barrier impairment, activation of intestinal immune response and increased frailty (7, 8). Therefore, there is a need to determine underlying mechanisms of host/microbiome age-related interactions, which may help identify interventions that could prevent dysbiosis and its negative effects on health. *Drosophila melanogaster* is an excellent model to study interactions between microbiota, intestinal aging, and organismal health. Increased microbial load in fly midguts has been associated with age-related intestinal dysfunction and increased mortality. It has been well documented that as flies age, bacterial load in the gut increases, contributing to pathology and, ultimately, death of the fly (7, 8, 10, 41, 43).

Here, we show that *Indy* reduction is associated with lower bacterial load and higher bacterial diversity in aging *Indy* flies compared to controls. We hypothesize that INDY as a plasma membrane citrate transporter affects microbiome via its impacts on cellular citrate levels. Reduction of SLC13A5 (the mammalian homolog of Indy) results in reduced intracellular citrate levels in mice (15). INDY has affinity for several other intermediate metabolites including succinate, malate, and fumarate, however, its highest affinity is for transporting citrate (12, 13). Another factor contributing to levels of Krebs cycle intermediates in the gut is the presence of microbes. Based on their genomes, *Acetobacter* is predicted to produce succinate while *Lactobacillus* likely consumes both succinate and citrate (46, 47), meaning that cross-feeding between gut microbes likely contributes to metabolite flux, resulting in downstream effects on nutrient cycling (48, 49). While citrate can promote growth of microbes such as *Lactobacillus*, it has also been shown to have bactericidal effects, possibly due to its ability to lower pH (50, 51). Thus, changes to citrate levels may result in alterations to diversity depending on how different bacteria are able to tolerate its effects. Recent study has shown that increased citrate consumption in young mice increases the SLC13A5 levels, increases bacterial count, decreases bacterial diversity and leads to increase colon permeability by affecting microbiome and activation of HIF-1α (52).

The mechanism by which age-associated increases in microbiome load decreases ISC homeostasis and contributes to shorter lifespan is thought to be tied to constitutive activation of the immune response, including release of reactive oxygen species and antimicrobial peptides, which damage enterocytes (7). This damage activates the JAK/STAT signaling pathway to repair and renew the gut epithelium, however, age-associated increases in bacterial load and dysbiosis are thought to lead to continuous JAK/STAT activation, resulting in hyperproliferation and loss of ISC homeostasis which causes further gut deterioration and dysplasia, and subsequently host death (42). We hypothesized that *Indy* reduction preserves midgut integrity and prevents intestinal barrier dysfunction by affecting host-microbiome interactions, thus preventing the negative effects of age-associated increase of JAK/STAT signaling pathway. In response to increased microbiome load with age, *Drosophila* cytokines Upd, Upd2 and Upd3 are released from different cell types and activate the JAK/STAT pathway. We found that *Indy* reduction prevented age-associated increases in microbiome load in male and female *Indy^206^*/+ flies compared to *yw* control at 7 days of age. We also found that *Indy* reduction increases microbiome diversity across the fly lifespan. This is consistent with our findings that *Indy* reduction decreases transcriptional levels of Upd2, and Upd3, but not other members of the JAK/STAT (Dome, 9STAT92E, Soc36E) pathway in the midgut of young *Indy* flies. Our data suggest that *Indy* reduction delays age-associated increase in the JAK-STAT signaling pathway, preventing its negative effect on ISC homeostasis. To examine the epistatic relationship between *Indy* and the JAK/STAT signaling pathway we determined lifespans of the *upd3^D^* mutant flies as homozygous, heterozygous and double heterozygous *upd3^Delta^*/+;*Indy^206^*/+. Homozygous *upd3^Delta^* flies have the shortest lifespans compared to heterozygous *upd3^Delta^*/+ and *Indy^206^*/+, which have similar and long lifespans. There is an additive effect on longevity when female flies are double heterozygous suggesting that the effect of *Indy* and *upd3* on female longevity partially overlaps. Our data suggest that *Indy* reduction has downstream effects on the microbiome of the fly, preventing bacterial overgrowth and altering microbiome diversity, which impacts longevity at least partially in a JAK/STAT-mediated fashion.

Age-related increases in ROS production contribute to activation of JAK/STAT pathway, which leads to ISC proliferation. We previously showed that *Indy* flies have reduced ROS production and increased transcription levels of genes encoding ROS-detoxification enzymes, which could reduce JAK/STAT activity (23). Another phenotype of *Indy* reduction in *D. melanogaster* is preserved mitochondrial function, which leads to reduced levels of reactive oxygen species in the fly (23). Genetic interventions that preserve mitochondrial capacity and lower ROS production, promote regenerative homeostasis in the *Drosophila* midgut (53–55). Recent studies have shown that transient exposure to low doses of oxidants in developing *D. melanogaster* results in reduction of *Acetobacter* spp. in the gut, which culminates in both extended gut healthspan and extended lifespan (56). In mice, mitochondrial function was shown to regulate reactive oxygen species production and, as a result, microbiome diversity (57), providing evidence that the effects of *Indy* reduction on the microbiome may have ties to its impacts on mitochondria and reactive oxygen species levels.

Despite the differences in the microbial communities of control flies and *Indy^206^/+*, our lifespan analysis comparing conventional and axenic flies make it clear that the microbiome is not required for *Indy*’s effects on longevity. However, given that total removal of the microbiome resulted in an even greater lifespan extension in *Indy^206^/+* compared to control flies over that observed in conventional flies, we expect that the microbiome does factor into *Indy*’s effects on the host. It is possible that the microbiome prevents *Indy^206^/+* flies aged in conventional conditions from reaching their maximum lifespan by, along with increasing ROS and JAK/STAT signaling, affecting upstream levels of citrate or other Krebs cycle intermediates, which translates to downstream effects of *Indy* on energy homeostasis (23). With the microbiome removed in axenic flies, these processes would be allowed to proceed unhindered, leading to a lifespan extension attributable to both the complete lack of microbiome and microbiome-independent factors. In summary, here we show that *Indy* reduction prevents age-associated increase in bacterial load and increases microbiome diversity, which reduce expression of Upd2 and Upd3 leading to delayed activation of the JAK/STAT signaling pathway. These changes together with preserved mitochondrial function and reduced ROS production reduce aberrant intestinal stem cell proliferation, the accumulation of mis-differentiated stem cells, and intestinal dysplasia, leading to healthier and longer life of *Indy* flies (Fig. 4N).

## Materials and Methods

### Fly stocks and rearing

The *yellow white (yw)* control line was obtained from the Bloomington Stock Center (Stock number 6599, *y[1[w[67c23]*). *Indy^206^* flies were originally obtained from Tim Tully (11, 58), *w[*]upd3]Delta]* was obtained from the Bloomington Stock Center (Stock number 55728). *yw* and *Indy^206^* flies were reared on food containing 25 mg/mL tetracycline for 3 generations to eliminate *Wolbachia*. This treatment was followed by growing flies for at least 10 generations in tetracycline-free food. Flies were recovered from antibiotics for at least six months prior to experiments to allow for regrowth of the microbiome. To addressed genetic background *Indy^206^* flies were backcrossed to *yw* flies for 10 generations prior experiments. Heterozygous *Indy^206^/+* flies were the F1 progeny of virgin *yw* female crossed with *Indy^206^/Indy^206^* male flies. Flies were reared on conventional food described in Supplemental Material. Stocks were maintained in a 25°C incubator with a 12-hour light:dark cycle, and newly emerged flies were passaged to fresh vials every 3-4 days.

### Generation of axenic flies

Generation of axenic flies is described in Supplemental Materials

### Lifespan Studies

Lifespan studies are described in Supplemental Materials.

### Culture-dependent microbiome analysis

Flies were surface sterilized in 70% ethanol then washed in sterile PBS. Individual whole flies were collected in 1.5 mL screw-top vials containing 1 mm glass beads and 500 µL of PBS. Samples were homogenized via bead-beating for 45 seconds at 4000 rpm. Homogenate was diluted to 10^-5^ in PBS using 96-well plates, then 3 µL of each dilution was pipetted onto MRS agar plates. Colony growth was recorded after 3-5 days. Colony types were determined visually, and colonies of different morphologies were re-streaked for isolation, then subjected to DNA extraction using the Epicentre MasterPure kit. Purified bacterial DNA was submitted to Eurofins for Sanger sequencing using primers 27F (5’-AGAGTTTGATYMTGGCTCAG-3’) and 519R (5’-GTATTACCGCGGCKGCTG-3’). DNA sequences were run through BLAST to obtain genus-level identification predictions (or species-level if possible).

### 16S rRNA microbiome sequencing

Flies were surface sterilized in 70% ethanol and washed in sterile PBS. Individual whole flies were collected in 1.5 mL screw-top vials containing a 1:1 mix of 1 mm glass beads and 0.5 mm glass beads and 300 µL of tissue and cell lysis buffer with proteinase K (Epicentre MasterPure DNA kit). Samples were homogenized via bead-beating for 45 seconds at 4000 rpm, then frozen at -80°C. Upon thawing, samples were again homogenized as performed previously to ensure tissue degradation. DNA extraction was performed following the Epicentre MasterPure protocol and 16S rRNA library preparation and sequencing was performed by the University of Connecticut Microbial Analysis, Resources, and Services (MARS) facility. Briefly, the V4 region of the 16S rRNA gene was PCR amplified using 515F and 806R primers with Illumina adapters and dual indices. Libraries were cleaned following the Omega Bio-Tek Mag-Bind Beads protocol and sequenced on the Illumina MiSeq (2x250 base pair kit).

### Bacterial culturing and generation of gnotobiotic flies

Bacterial strains isolated from the culture-based microbiome analysis were kept at -80°C in 50% glycerol for long-term storage. To generate gnotobiotes, bacteria were streaked on MRS agar plates, then single colonies were transferred to flasks containing a 1:1 mix of MRS broth and mannitol broth. Liquid cultures were grown shaking at 29°C, 200 rpm, for up to 24 hours. Cultures were set to an OD_600_ of 0.5 (about 10^5^ cells per mL), and 150 mL of a 1:1 mix of diluted culture and 2.5% sucrose was fed to axenic adults. After several days of laying, adults were discarded and F1 progeny were collected for lifespan analysis.

### Bioinformatics and statistical analyses

16S rRNA sequences were processed using Mothur v. 1.39.4 following the MiSeq SOP (59). Sequences were clustered at 97% similarity and rarified to 10,000 reads for alpha and beta diversity metrics. Taxonomy plots, Shannon alpha diversity plots, and Bray-Curtis beta diversity principal component analysis plots were generated in R using ggplot2. Lifespan was analyzed in GraphPad Prism using Kaplan-Meier survival; statistical significance of differences in survival between genotypes was determined via Log-rank test. Statistical differences between *yw* controls and *Indy* heterozygote CFU counts were determined via two-way ANOVA using GraphPad Prism. Significance is expressed as: P>0.05=ns, P£0.05=*, P£0.01=**, P£0.001=***, P£0.0001=****.

### RNA-Seq and Analysis (RiboZ-Seq)

For RNA-Seq experiments flies were collected and maintained as described above. At ages 7 and 40, *Indy^206^* heterozygous and *yw* flies, aged on conventional and axenic condition were separated by sex on CO_2_ and midguts were dissected from female flies. Total RNA was isolated from 3 biological replicates from 30-40 female flies in each replicate, except samples from 40 days old *yw* female aged in axenic condition, which had between 17 and 26 guts in each replicate. Total RNA was isolated using Trizol as described (23). One μg of each RNA sample was used to prepare RNA-seq libraries with the Illumina Tru-Seq Stranded mRNA Library Preparation kit (cat # 20020594) with the IDT for Illumina TruSeq RNA UD Indexes set (cat # 20040871). The libraries were pooled and sequenced on one lane of a NovaSeq 6000 S4 flow cell generating 100bp paired-end reads yielding an average of 80 million reads per sample and a minimum of 20 million reads per replicate.

### mRNA-Seq analysis

The quality of the sequencing data was evaluated with FastQC (v0.11.9) and MultiQC (1.9). Sequence reads were mapped to the *Drosophila* genome (GSE97233) with STAR (version) (2.71a) and the resulting BAM files were used to generate gene counts with featureCount using the uniq-counting mode. Analyses of sample relationship and differential expression were performed with DESeq2 (60). To identify specific pathways associated with diet shift, we performed functional enrichment analysis using the DAVID database. Pathway analysis and identification of networks associated with shifting flies were also carried out by Gene Ontology (GO) and the KEGG.

## Author Contributions

Conceptualization, B.R., N.A.B; Methodology and Investigation, D.N.A.L., S.P, J.M., E.S., K.K., S.O., B.R.G., B.R., N.A.B; Writing – Original Draft and Figure Creation, D.N.A.L., E.S., K.K., S.O., B.R., N.A.B; Writing – Review & Editing, Supervision, D.N.A.L., E.S., S.O., B.R., N.A.B

## Classification

Biological Sciences, Genetics/Physiology.

## Data sharing plans

All the bulk RNA-Seq data have been deposited at NCBI BioProject and are publicly available as of the date of publication. Submission ID: SUB16031777 BioProject ID: PRJNA1431527. Raw data for survival and microbiome analysis (CFUs, 16S rRNA pipeline) is available in supplemental data and posted on Figshare (DOI to be updated upon publication).

## Acknowledgments

This work was supported by grants from the National Institute of Health: R35GM128871 to NAB; RO1AG059586, R01AG059586-03S1, R56AG082788, the University of Connecticut (UConn) Claude D. Pepper Older Americans Independence Center (P30-AG067988) to B.R.; U24 HG009889, and R35 GM118140 to BRG; Rogina is a recipient of a Glenn Award for Research in Biological Mechanisms of Aging. Figures 2A and 4O were prepared using BioRender.

## Supporting Information

### Supplementary Materials

#### Fly Food Preparation

Flies were reared on food containing per liter of water: 28 grams Brewer’s yeast, 49 grams yellow cornmeal, 113 grams sucrose, 8.1 grams *Drosophila* agar, and 2.4 grams Tegosept (Methyl4-hydroxybenzoate) dissolved in 10.7 mL of 100% ethanol as in (1). Food was autoclaved prior to addition of Tegosept.

#### Generation of axenic flies

Using embryo cages, *yw* and *Indy^206^/Indy^206^* flies laid eggs overnight on grape juice agar plates smeared with yeast paste. The next morning, plates were rinsed with 70% ethanol, then PBS, and embryos were lifted from the agar with sterile swabs. Embryos were poured into cell strainers and rinsed with 5% bleach until dechorionation was confirmed under dissecting microscope. Embryos were then rinsed well with sterile water and suspended in 100% ethanol before being transferred via pipette to sterile fly food to develop. Axenic flies were maintained by passaging flies only in a biosafety cabinet or next to a flame. To mate axenic *yw* and *Indy^206^/Indy^206^* flies or collect axenic flies for experiments, CO_2_ fly pad and paintbrush were thoroughly bleached, then treated with 70% ethanol, then put under UV light for 15 minutes in a PCR cabinet, where flies were then collected. Stocks and experimental vials containing axenic flies were regularly screened visually and via plating to ensure axenic status was maintained.

#### Lifespan Studies

Newly emerged *yw* or F1 *Indy^206^/+* (cross described in Fly Stock Rearing) flies were collected and passed onto fresh food twice per week (Fig. 1A-D) or daily (Fig. 1E,F). Each vial contained 20 male and 20 female flies, and 10 replicate vials per treatment were established. Deceased flies were recorded regularly until all flies in each vial were dead. Heterozygous flies used in survivorship analysis described in Fig. 5H,I were: a) *Indy^206^*/+ were F1 generation from crosses in which virgin *Indy^206^/Indy^206^* homozygous female flies were crosses to *yw* males; b) *upd3^D^*/+ were F1 generation of cross in which virgin *upd3^D^/upd3^D^*homozygous female flies were crossed to *yw* males; c) *upd3^D^*/+; *Indy^206^*/+ were F1 generation from crosses in which virgin *upd3^D^/upd3^D^*homozygous female were crossed to *Indy^206^/Indy^206^*homozygous male flies. F1 progeny for lifespan studies were collected within 24 hours following eclosion using CO_2_ and transferred to a vial at density of 25 male and 25 female flies per vial containing conventional corn diet. 12 replicate vials were established. Flies were maintained in a humidified temperature-controlled environmental chamber at 25°C (Percival Scientific) on a 12-hour light:dark cycle. Flies were passed every day, and the number of dead flies were counted. Longevity data were censored for early mortality (1-10 Days) to remove death due to post-eclosion maturation, or other deaths that are not related to aging. The number of censored flies is listed in brackets in Tables S1, S2, and 5. Survivorship curves were analyzed by log-rank test JMP16 program.

**Fig. S1.**
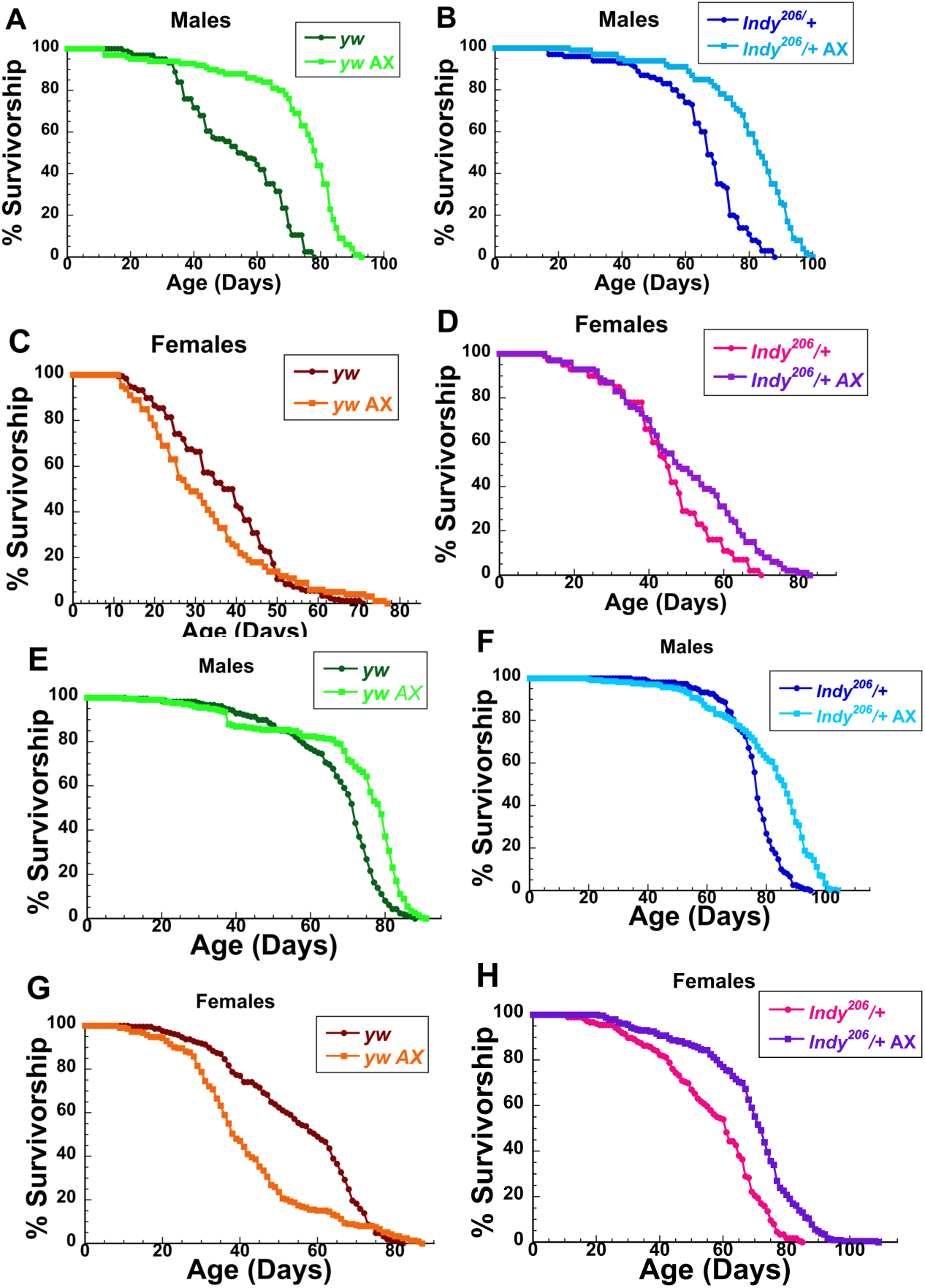
Effects of axenic culture on lifespan of control and *Indy^206^/+* flies passed twice per week (A-D) or daily (E-H). Axenic conditions extend lifespan of *yw* (A,E) and *Indy^206^/+* (B,F) males, and *Indy^206^/+* females (D,E), but *yw* females live shorter in axenic condition (C,G). Longevity of conventional (green) and axenic (light green) *yw* males (A,E); conventional (brown) and axenic (orange) *yw* females (C,G); conventional (blue) and axenic (light blue) *Indy^206^/+* males (B,F); conventional (magenta) and axenic purple *Indy^206^/+* female (D,H) flies expressed as Kaplan-Meier survival curves. N= 131-187 individual flies per condition spread across 10 vials that were passaged twice weekly. AX: axenic

**Fig. S2.**
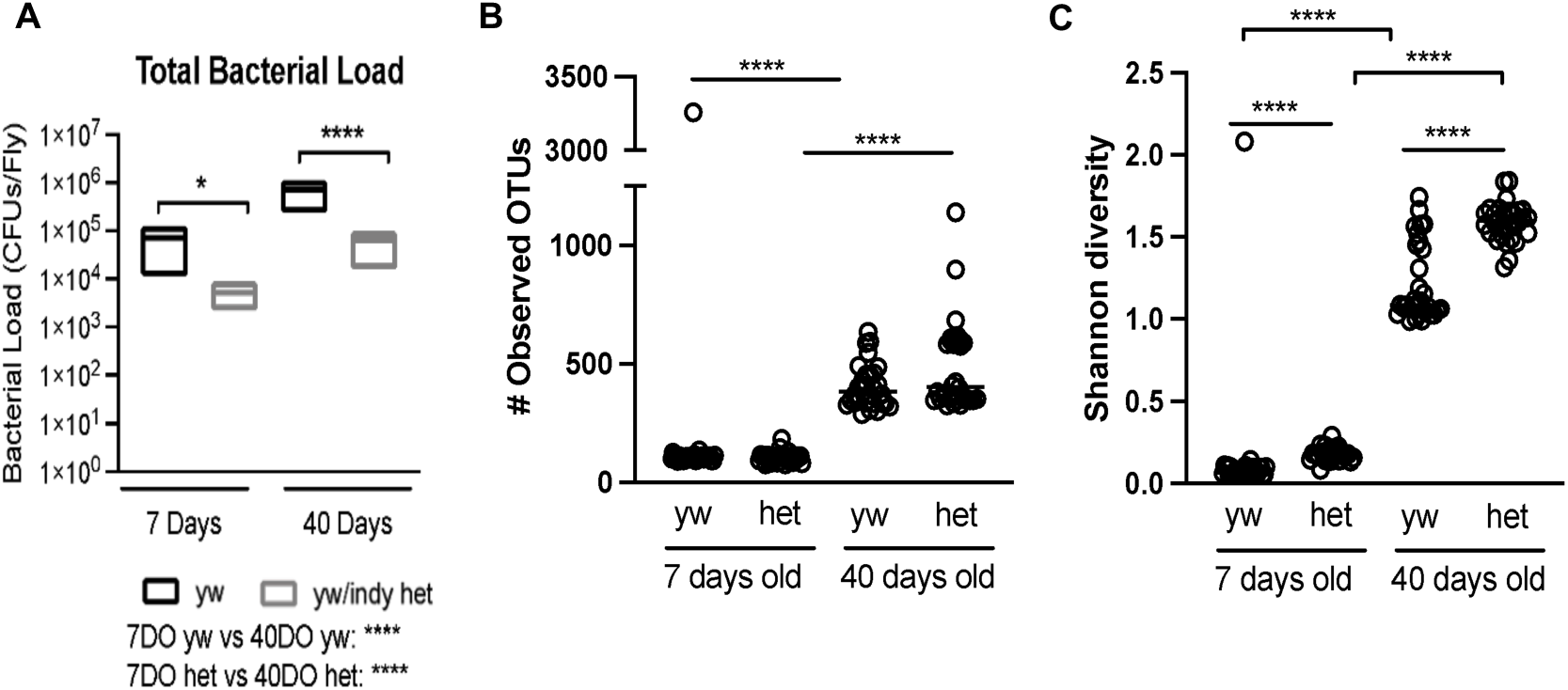
Bacterial load and diversity in young and old controls and *Indy^206^/+* heterozygous flies. **A)** Total bacterial load in 7-and 40-day old *yw* control and *Indy^206^/+* female flies. Bacterial counts are expressed as colony forming units (CFU) per fly, as determined by growth of colonies on MRS agar. Lines and error bars represent median and interquartile range of data points. Statistical differences between *yw* control and *Indy^206^/+* female flies CFU counts were determined via two-way ANOVA. N= 3 replicate experiments collecting 3 individual flies per treatment group (n=9 flies per genotype/age). Statistical results not shown were non-significant. Significance is expressed as follows: ns, not significant (*P* > 0.05); *, *P* ≤ 0.05; **, *P* ≤ 0.01; ***, *P* ≤ 0.001; ****, *P* ≤ 0.0001. **B)** Alpha diversity of young and old control, and *Indy^206^/+* heterozygous female flies determined by sequencing V3V4 region to better reach species level IDs.

**Fig. S3:**
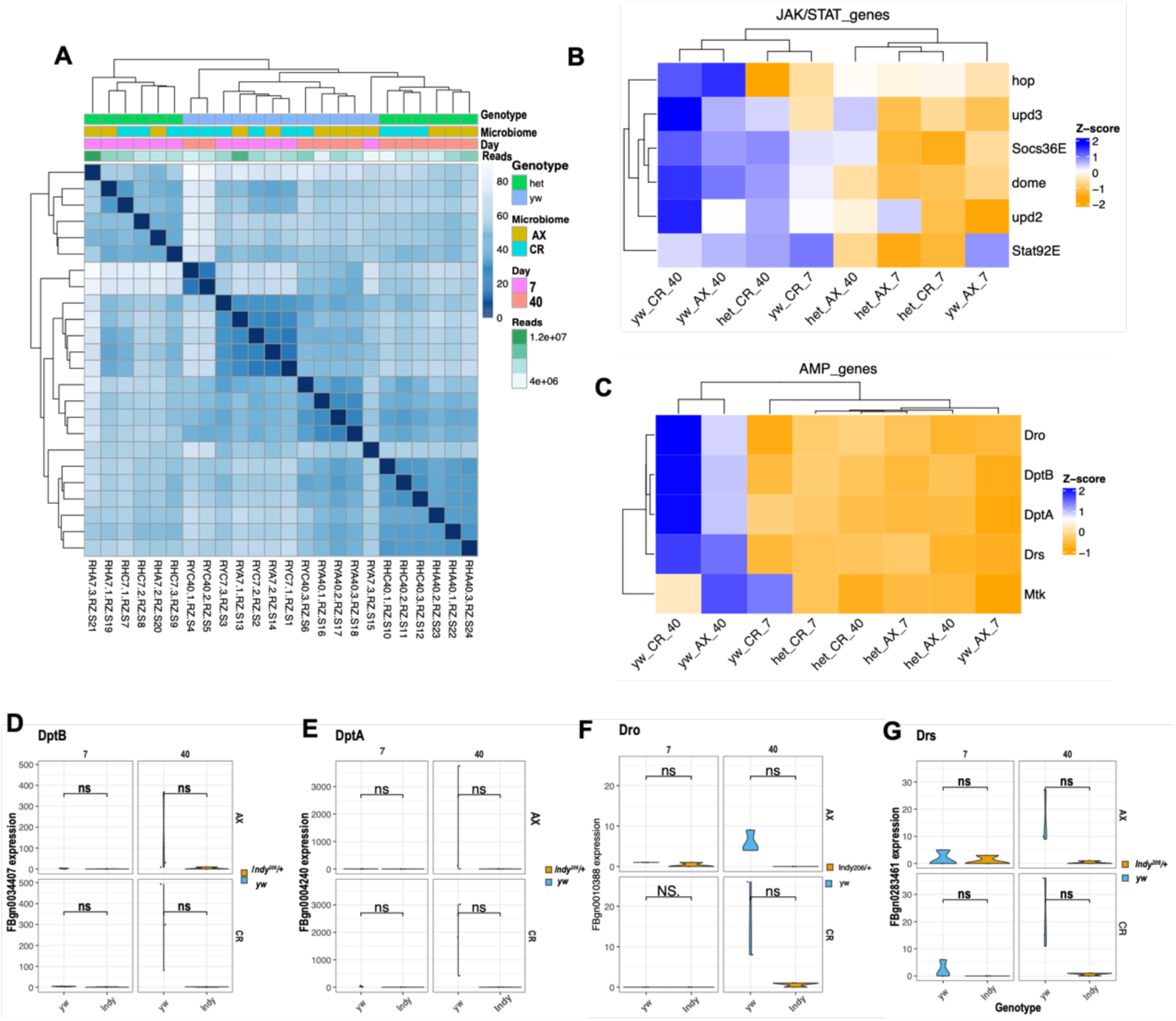
**A)** A sample-to-sample distance heatmap shows the driving force for distance in each genotype (*yw*: control, het: *Indy^206^/+)*, age (7, 40 days) and rearing condition (CD: conventional, AX: axenic condition). **B)** Heatmaps of members of JAK/STAT signaling pathway: Upd3, Upd2, Hop, Dome, Socs36E and Stat92E, in control and *Indy^206^/+* flies at 7 and 40 days of age in conventional and AX conditions. **C)** Heatmaps of members of antimicrobial peptide (AMPs): Diptericin A (Dpt A), Diptericin B (Dpt B), Drosocin (Dro), and Drosomycin (Drs), in control and *Indy^206^/+* flies at 7 and 40 days of age in conventional and AX conditions. **D-G)** The levels of DptB, DptA, Dro and Drs are not different in the midgut of *Indy* and control female flies.

**Table S1.**
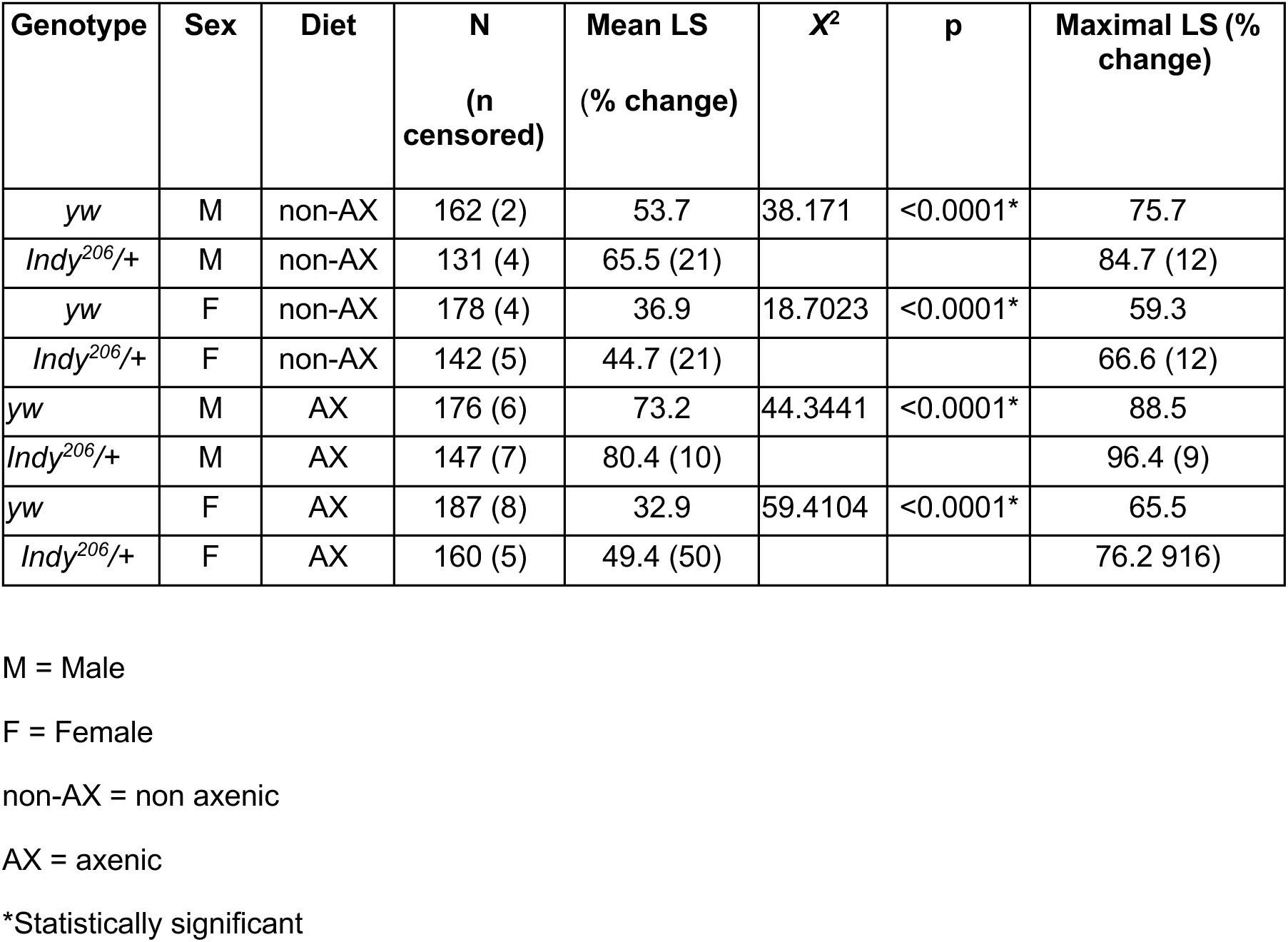
Effects of reducing *Indy* gene activity on longevity of flies aged in non-axenic or axenic condition compared to control *yw* flies when passed twice per week.

**Table S2:** Effects of axenic culture on lifespan of control and *Indy^206^/+* flies passed twice per week.

| Genotype | Sex | Diet | N<br>(n<br>censored) | Mean<br>LS<br>(%<br>change) | $\chi^2$ | p | Maximal LS (%<br>change) |
| --- | --- | --- | --- | --- | --- | --- | --- |
| *yw* | M | non-AX | 162 (2) | 53.7 | -<br>181.8958 | <0.0001* | 75.7 |
| *yw* | M | AX | 176 (6) | 73.2 (36) |  |  | 88.5 (17) |
| *yw* | F | non-AX | 178 (4) | 36.9 | 2.5089 | 0.1132 | 59.3 |
| *yw* | F | AX | 187 (8) | 32.9 (-10) |  |  | 65.5 (10) |
| *Indy*<sup>206/+</sup> | M | non-AX | 131 (4) | 65.5 | 106.5593 | <0.0001* | 84.7 |
| *Indy*<sup>206/+</sup> | M | AX | 147 (7) | 80.4 (23) |  |  | 96.4 (14) |
| *Indy*<sup>206/+</sup> | F | non-AX | 142 (5) | 44.7 | 14.7471 | <0.0001* | 66.6 |
| *Indy*<sup>206/+</sup> | F | AX | 160 (5) | 49.4 (11) |  |  | 76.2 (14) |

non-AX = non-axenic
AX = axenic
\*Statistically significant

**Table S3:** Effects of reducing *Indy* gene activity on lifespan of control and *Indy^206^/+* flies aged in non-axenic or axenic condition when passed daily.

| Genotype | Sex | Diet | N<br>(n censored) | Mean LS<br>(% change) | X<sup>2</sup> | p | Maximal LS (%<br>change) |
| --- | --- | --- | --- | --- | --- | --- | --- |
| *yw* | M | non-AX | 208 (0) | 67.2 | 49.9147 | <0.0001* | 82.7 |
| *Indy*<sup>206/+</sup> | M | non-AX | 149 (1) | 75.7 (13) |  |  | 89.3 (8) |
| *yw* | F | non-AX | 208 (3) | 55.3 | 1.7025 | 0.1806 | 75.5 |
| *Indy*<sup>206/+</sup> | F | non-AX | 176 (5) | 56.7(3) |  |  | 77.7 (3) |
| *yw* | M | AX | 198 (3) | 72.1 | 94.1875 | <0.0001* | 87.0 |
| *Indy*<sup>206/+</sup> | M | AX | 208 (2) | 81.0 (12) |  |  | 99.8 (6) |
| *yw* | F | AX | 212 (3) | 42.8 | 169.5708 | <0.0001* | 76.9 |
| *Indy*<sup>206/+</sup> | F | AX | 230 (3) | 68.8 (61) |  |  | 91.5 (21) |

M = Male F = Female non-AX = non-axenic AX axenic \*Statistically significant

**Table S4:** Effects of axenic culture on lifespan of control and *Indy^206^/+* flies when passaged daily.

| Genotype | Sex | Diet | N | Mean LS<br>(% change) | X<sup>2</sup> | p | Maximal LS (%<br>change) |
| --- | --- | --- | --- | --- | --- | --- | --- |
| *yw* | M | non-AX | 208 (0) | 67.2 | 56.8646 | <0.0001* | 82.7 |
| *yw* | M | AX | 198 (3) | 72.1 (7) |  |  | 87.0 (5) |
| *yw* | F | non-AX | 208 (3) | 55.3 | 27.0522 | <0.0001* | 75.5 |
| *yw* | F | AX | 212 (3) | 42.5 (-23) |  |  | 76.9 (2) |
| *Indy*<sup>206/+</sup> | M | non-AX | 149 (1) | 75.7 | 68.4399 | <0.0001* | 89.3 |
| *Indy*<sup>206/+</sup> | M | AX | 208 (2) | 81.0 (7) |  |  | 99.7 (12) |
| *Indy*<sup>206/+</sup> | F | non-AX | 176 (5) | 56.7 | 73.5511 | <0.0001* | 77.7 |
| *Indy*<sup>206/+</sup> | F | AX | 230 (3) | 68.8 (21) |  |  | 91.5 (18) |

M = Male
F = Female
non-AX = non-axenic
AX = axenic
\*Statistically significant

**Table S5:**
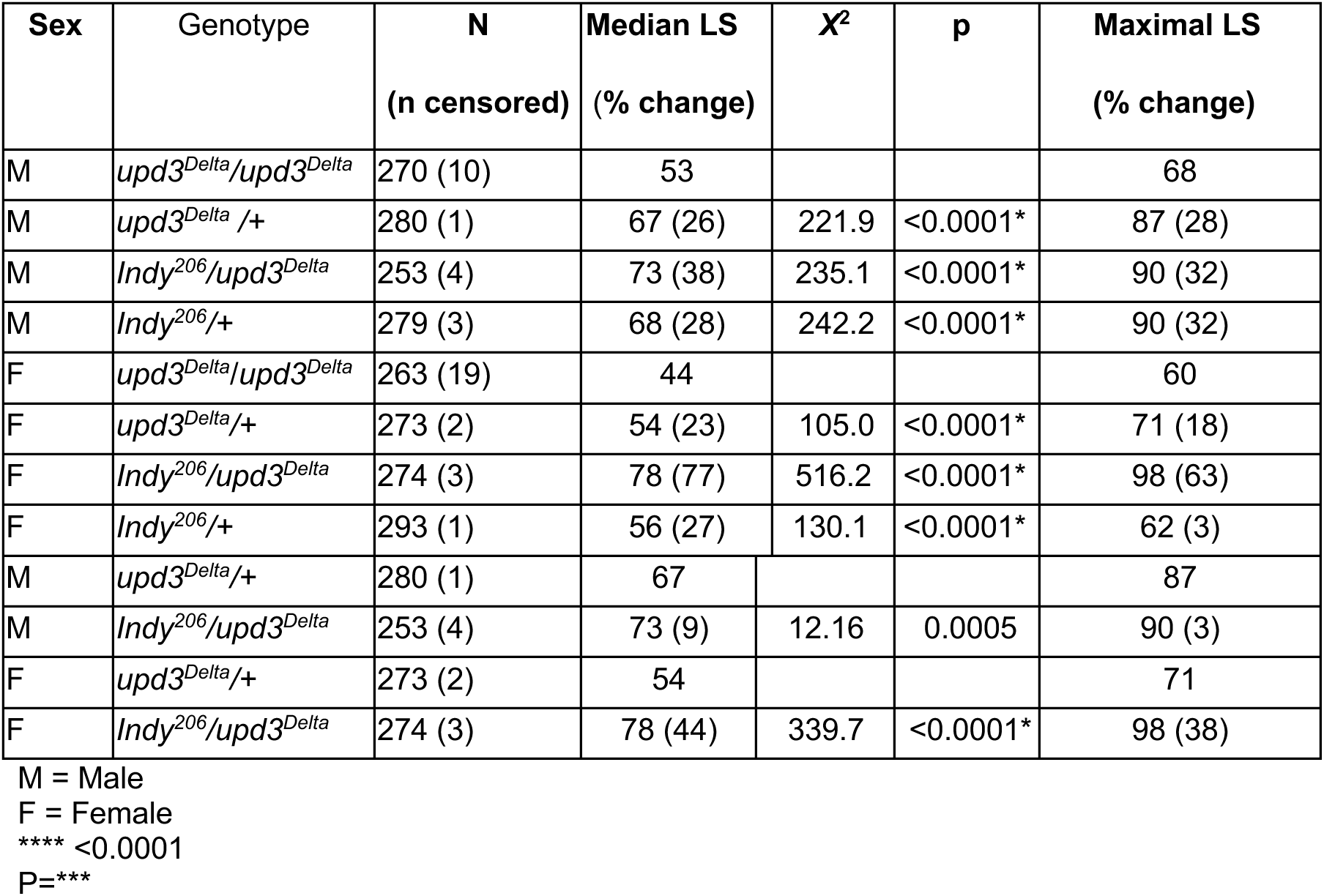
Effect of reducing *Indy* gene activity on lifespan of *upd3^Delta^* flies.

| Sex | Genotype | N<br>(n censored) | Median LS<br>(% change) | X<sup>2</sup> | p | Maximal LS<br>(% change) |
| --- | --- | --- | --- | --- | --- | --- |
| M | *upd3<sup>Delta</sup>/upd3<sup>Delta</sup>* | 270 (10) | 53 |  |  | 68 |
| M | *upd3<sup>Delta</sup>/+* | 280 (1) | 67 (26) | 221.9 | <0.0001* | 87 (28) |
| M | *Indy<sup>206</sup>/upd3<sup>Delta</sup>* | 253 (4) | 73 (38) | 235.1 | <0.0001* | 90 (32) |
| M | *Indy<sup>206</sup>/+* | 279 (3) | 68 (28) | 242.2 | <0.0001* | 90 (32) |
| F | *upd3<sup>Delta</sup>/upd3<sup>Delta</sup>* | 263 (19) | 44 |  |  | 60 |
| F | *upd3<sup>Delta</sup>/+* | 273 (2) | 54 (23) | 105.0 | <0.0001* | 71 (18) |
| F | *Indy<sup>206</sup>/upd3<sup>Delta</sup>* | 274 (3) | 78 (77) | 516.2 | <0.0001* | 98 (63) |
| F | *Indy<sup>206</sup>/+* | 293 (1) | 56 (27) | 130.1 | <0.0001* | 62 (3) |
| M | *upd3<sup>Delta</sup>/+* | 280 (1) | 67 |  |  | 87 |
| M | *Indy<sup>206</sup>/upd3<sup>Delta</sup>* | 253 (4) | 73 (9) | 12.16 | 0.0005 | 90 (3) |
| F | *upd3<sup>Delta</sup>/+* | 273 (2) | 54 |  |  | 71 |
| F | *Indy<sup>206</sup>/upd3<sup>Delta</sup>* | 274 (3) | 78 (44) | 339.7 | <0.0001* | 98 (38) |

M = Male F = Female \*\*\*\* <0.0001 P=\*\*\*

